# E2F-dependent genetic oscillators control endoreplication

**DOI:** 10.1101/858746

**Authors:** Minhee Kim, Nam-Sung Moon

## Abstract

Polyploidy is an integral part of development and is associated with cellular stress, aging and pathological conditions. The endoreplication cycle, comprised of successive alternations of G and S phases without cell division, is widely employed to produce polyploid cells. The endocycle is driven by continuous oscillations of Cyclin E/Cdk2 activity, which is governed by E2F transcription factors. In this study, we provide mechanistic insight on how E2F-dependent Cdk oscillations during endocycles are maintained in Drosophila salivary glands. Genetic experiments revealed that an alternative splicing isoform of E2F1, E2F1b, regulates the circuitry of timely S phase entry and exit by activating a subset of E2F target genes. E2F1b regulates the Drosophila ortholog of p27^CIP/KIP^-like Cdk inhibitor Dacapo to precisely time S phase entry by controlling the CycE/Cdk2 activity threshold. Upon entry to S phase, E2F1b-dependent PCNA expression establishes a negative feedback loop through the PIP box-mediated degradation of E2F1. Overall, our study uncovers a network of E2F-dependent genetic oscillators that are critical for the periodic transition between G and S phases during endoreplication.

## Introduction

The endoreplication cycle (endocycle) is a variant cell cycle comprised of successive alternations of G and S phases, leading to increased DNA content without cell division [1]. This alternate cell cycle is widely used to produce polyploid cells during development and for maintaining tissue homeostasis [2]. Endoreplication is driven by oscillations of Cyclin-dependent kinase (Cdk) activity, where low Cdk activity promotes pre-replicative complex (pre-RC) formation and high Cdk activity initiates S phase [3]. At the center of regulating Cdk activity are E2F transcription factors. For example, E2F7 and 8 coordinate the balance between endoreplication and the mitotic cycle during development of placental trophoblast giant cells and hepatocytes of the liver [4,5]. In Drosophila, specific defects in endoreplicating tissues were observed in three independent viable alleles of the only activator E2F, *de2f1: de2f1^i2^, de2f1^su89^, and de2f1b* [6–9]. Interestingly, while mitotic tissues develop with no apparent defects, defects in endoreplicating tissues, such as the salivary gland, ovary, and bristles have been described in these alleles. Overall, studies in various organisms have demonstrated the importance of E2Fs for timely expression of S phase cyclins [10] and suppressing expression of mitotic cyclins during the endocycle [4,5].

The relationship between E2F and S phase cyclin, Cyclin E (CycE), is well-characterized in the Drosophila salivary gland. E2F1 accumulation during G phase and its subsequent degradation during S phase ensures timely expression of CycE [11]. Degradation of E2F1 is carried out by CRL4^Cdt2^, an E3 ubiquitin ligase, which targets E2F1 via the Proliferating Cell Nuclear Antigen (PCNA) Interacting Peptide (PIP) degron in a DNA replication-dependent manner [12,13]. While the PIP degron-dependent mechanism seems to be unique to Drosophila, limiting E2F activity upon entry to S phase in endocycling tissues seems to be common and essential across numerous organisms [11]. For example, one of the crucial targets of E2F7/8 is mammalian E2F1[4,5,14,15]. Recently, we reported that an alternatively spliced isoform of the activator *de2f1* named E2F1b is critical for endocycling tissues, such as the salivary gland [9]. In this study, we further investigated the precise role of E2F1b during endoreplication in the salivary gland. Our findings reveal that E2F1b establishes feedback circuits that are necessary for proper entry to and exit from S phase. In G phase, E2F1b-dependent activation of Dacapo (Dap) limits CycE/Cdk2 activity. This likely ensures that appropriate levels of factors necessary for proper DNA synthesis are achieved prior to entering S phase. In S phase, E2F1b-dependent expression of PCNA helps establish a negative feedback loop that targets E2F1 for degradation during S phase, directly linking E2F1 activity to its own destruction. Overall, our study demonstrates that E2F1b-specific targets establish the genetic network that sets up the oscillating nature of G and S phase during endocycle progression.

## Results and Discussion

### E2F1b is required for biphasic CycE and E2F1 oscillations in endocycling salivary glands

To understand the impact of E2F1b deregulation during the endocycle, we carefully analyzed actively cycling salivary glands, staged 80-85 hours After Egg Laying (hr AEL). Previous studies have demonstrated that the biphasic expression of core molecular oscillators, E2F1 and CycE, is required for endocycle progression [13]. Indeed, this biphasic expression is clearly visible when a contiguous plot profile of salivary gland cells marked with E2F1 and CycE along with its associated 2-dimensional (2D) scatter plot are generated (Figure 1A). As alluded from our previous study, the biphasic oscillation of CycE and E2F1 expression is disrupted in *de2f1b* salivary glands ([9] and Figure 1B). The relationship between CycE and E2F1 is no longer mutually exclusive and rather becomes linear, indicating that E2F1b is required for the periodic nature of CycE and E2F1 expression during endocycles. Importantly, E2F1a is readily detected in the *de2f1b* salivary gland, indicating that this E2F1b’s function during endoreplication cannot be compensate by E2F1a (Figure 1B).

**Figure 1.**
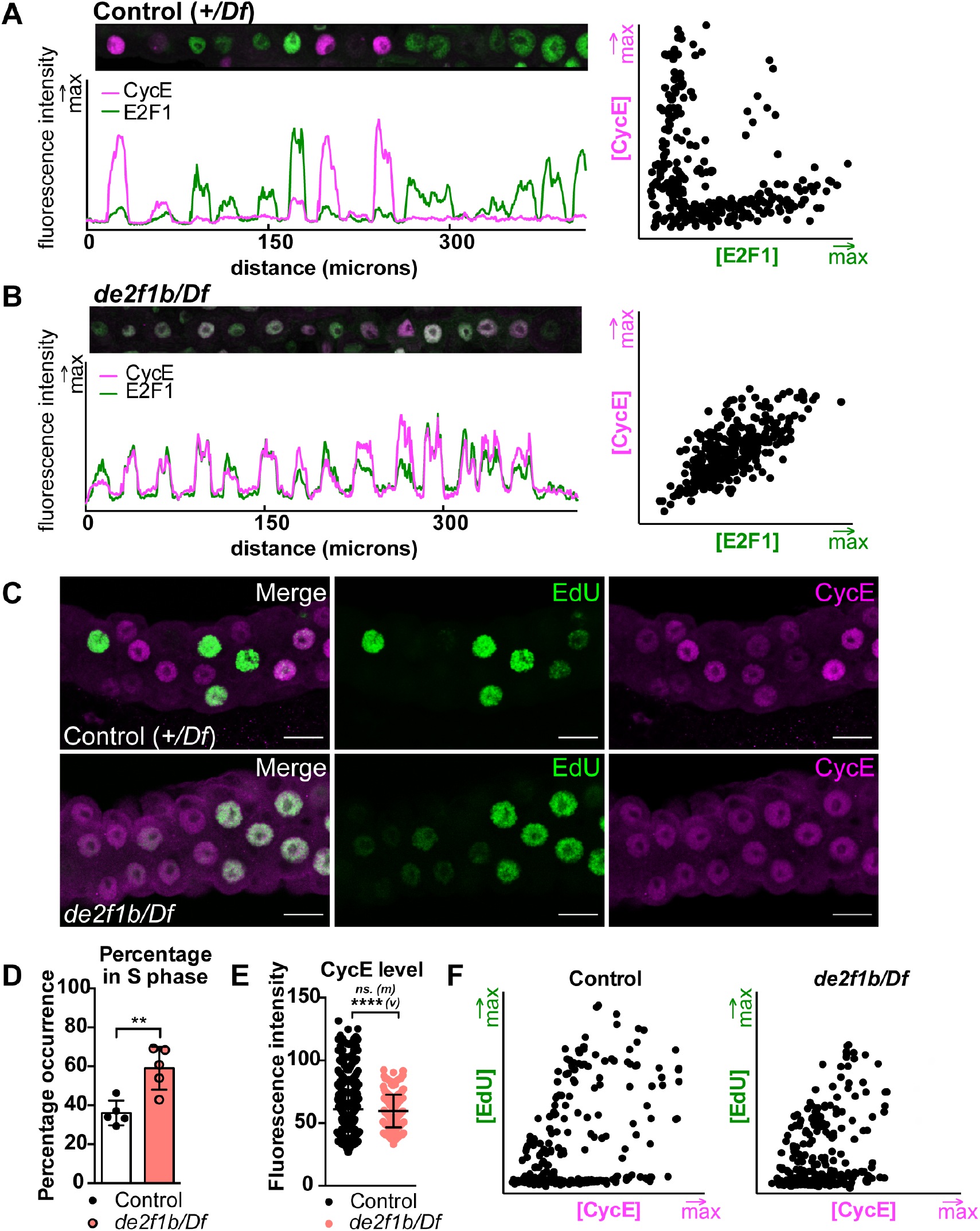
E2F1b is required for biphasic CycE and E2F1 oscillations in endocycling salivary glands. (A, B) A plot profile (left) and 2D scatter plot (right) depicting the nuclear intensities of E2F1 (green) and CycE (magenta) from 80-85 hours after egg laying (AEL) salivary gland cells in Control (A) and *de2f1b/Df* (B) backgrounds. Confocal micrographs of 80-85 hr AEL salivary glands marking EdU (green) and CycE (magenta) of indicated genotypes. Scale bars = 25 μm. (D) The percentage of EdU positive cells are quantified (n=5/genotype). Error bars indicate standard deviation (s.d.). ** = p ≤ 0.01. (E) A scatter plot depicting CycE protein intensity where the mean (m) CycE is not significantly different but the variance (v) is significantly changed between the two genotypes. ns = p > 0.05; **** = p ≤ 0.0001. (F) 2D scatter plots showing intensities of CycE and EdU between control and *de2f1b/Df*.

To determine the impact of disrupted E2F1 and CycE expression during endocycle, ethynyl deoxyuridine (EdU) incorporating S phase cells relative to CycE expression was determined. Interestingly, despite the lack of mutual exclusivity between E2F1 and CycE, endocycle progresses in *de2f1b* salivary glands, evident by the abundance of EdU positive cells (Figure 1C). In fact, the percentage of S-phase cells in *de2f1b* salivary glands is significantly higher than in the control (Figure 1D). Notably, quantification of overall CycE intensity revealed that although the mean CycE expression level between the control and *de2f1b* salivary glands is not significantly different, cells with high CycE expression are lacking in *de2f1b* salivary glands (Figure 1E). Consequently, the overall variance of CycE expression among different nuclei is significantly decreased in *de2f1b* mutants (Figure 1D). Comparing 2D scatter plots of CycE and EdU intensities of each nuclei showed that the overall pattern of the 2D scatter plot is not changed. This suggests that while CycE high cells may be scarce, the relationship between CycE expression and EdU incorporation is not altered in *de2f1b* salivary glands (Figure 1F). Taken together, these data suggest that while E2F1b is required for establishing the oscillation of E2F1 and CycE expression, *de2f1b* mutant cells readily enter S phase during the endocycle.

### Cells in *de2f1b* salivary glands prematurely enter S phase

Next, we used the Fly-Fluorescent Ubiquitination-based Cell Cycle Indicator (FUCCI) system to better characterize cell cycle-dependent events in *de2f1b* salivary glands [16]. Specifically, the G1 sensor, hereon referred to as G1-FUCCI, was used to positively mark G phase cells. G1-FUCCI is GFP fused to PIP box-dependent E2F1 degron (GFP-E2F1_1-230_), which is targeted for degradation in a S phase-specific manner (Figure 2A) [16,17]. To unambiguously mark G and S phase cells, G1-FUCCI and EdU incorporation was used (Figure 2B). 2D scatter plot comparing GFP and EdU intensities from control salivary glands showed that the population of cells with G1-FUCCI expression are distinct from EdU incorporating cells (Figure 2C and 2D left panels). Notably, the distinction between G1-FUCCI and EdU populations are lost in *de2f1b* salivary glands (Figure 2C and 2D right panels), suggesting that S phase-specific PIP box-dependent degradation mechanism is deregulated in *de2f1b* salivary glands. This likely explains the reduced oscillation and mutually exclusiveness between E2F1 and CycE (Figure 1A and 1B) and also suggests that the boundary between G and S phases becomes unclear in *de2f1b* salivary glands (Figure 2D bottom panel).

**Figure 2.**
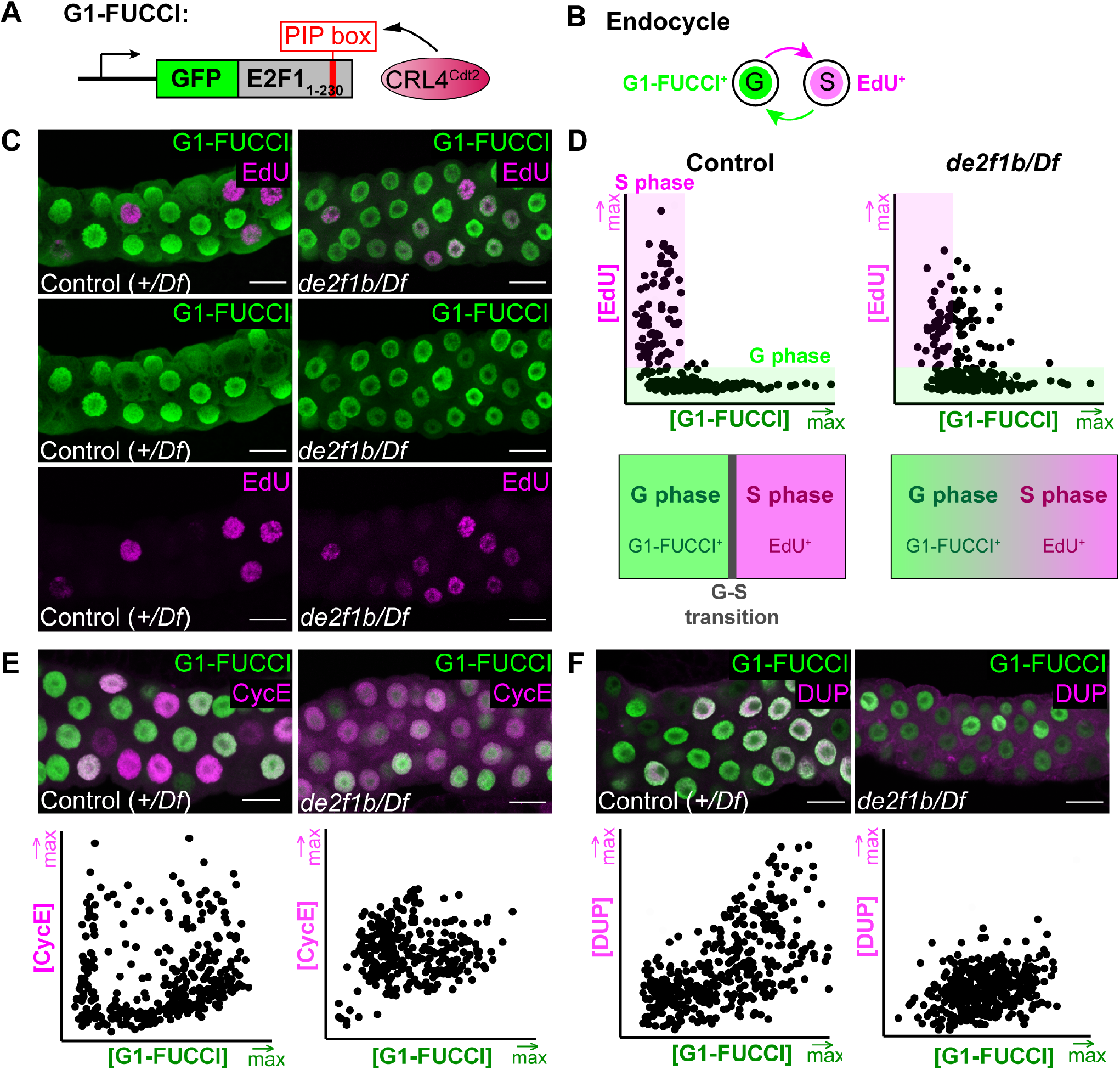
Cells in *de2f1b* salivary glands prematurely enter S phase. (A) A schematic of G1-FUCCI, which contains the PIP box of E2F1 targeted by CRL4^Cdt2^. (B) A schematic describing the use of G1-FUCCI and EdU to visualize G and S phase cells, respectively. (C) Confocal micrographs of 80-85 hr AEL salivary glands of control and *de2f1b.* G and S phase cells are visualized via G1-FUCCI and EdU, respectively. (D) 2D scatter plots depicting G1-FUCCI and EdU intensities of control and *de2f1b* salivary glands. The analysis indicates that the G-S phase boundary in *de2f1b* salivary glands may not be as precise as control (Schematics in lower panel). (E) Confocal micrographs of 80-85 hr AEL salivary glands of control and *de2f1b*. Relative expression pattern of CycE to G1-FUCCI is determined and 2D scatter plots depicting their expression pattern is presented in the lower panel. (F) The experiment described in E is performed to determine the relative expression pattern of Double-Parked (DUP) to G1-FUCCI. All scale bars = 25 μm.

To understand how *de2f1b* cells readily enter S phase without accumulating high levels of CycE, we determined the expression pattern of CycE relative to G1-FUCCI. Notably, in addition to G1-FUCCI negative cells (S phase cells) expressing high CycE, there is a large fraction of G1-FUCCI positive cells also expressing high CycE (Figure 2E left panel). This observation shows that CycE can accumulate at high levels during G phase and that its levels may not directly reflect its activity. This also suggests that there may be a mechanism in place to prevent CycE/Cdk2 activity during G phase. We hypothesized that this pool of G phase cells likely represents late G phase cells that are primed to enter S phase, accumulating high level of CycE prior to being activated in S phase. While CycE high cells are scarce in *de2f1b* salivary glands, they are generally associated with lower levels of G1-FUCCI and lacking from high G1-FUCCI population (Figure 2E right panel). This result indicates that the CycE high G phase population observed in the control is missing in *de2f1b* salivary glands. This result, together with the observation that cells in *de2f1b* salivary glands readily enter S phase (Figure 1E), suggests that the loss of E2F1b allows cells to enter S phase without accumulating high levels of CycE. To further test this idea, we compared the expression of Double Parked (DUP), a pre-RC component also known as Cdt1 [18], and G1-FUCCI. DUP accumulates in G phase where Cdk activity is low and rapidly degraded at the onset of S phase [19–22]. Consistent with this idea, DUP demonstrates a linear relationship with G1-FUCCI in control salivary glands, DUP is low where G1-FUCCI is low and DUP is high where G1-FUCCI is high (Figure 2F left panel). Strikingly, in *de2f1b* salivary glands, DUP high G phase population is absent, again indicating a failure to accumulate appropriate DUP levels prior to entering S phase (Figure 2F right panel). Taken together, our results suggest that *de2f1b* cells fail to accumulate specific factors in G phase and prematurely enter S phase during endocycle.

### Specificity of E2F1b-dependent transcription during the endocycle

To gain mechanistic insights behind the endocycle defect, we determined the expression levels of E2F targets in *de2f1b* salivary glands. E2F1 transcriptionally regulates expression of core endocycle machineries, such as *cycE* as well as genes important in mitotic cycles [13]. Reverse transcriptase-quantitative polymerase chain reaction (RT-qPCR) was performed on RNA collected from 80-85 hr AEL salivary glands of control and *de2f1b*. We selected E2F targets that were specific to G1/S or G2/M to determine if a specific group of E2F1 target is regulated by E2F1b. As shown in Figure 3A, E2F1b deregulation specifically affects G1/S targets and not G2/M targets.

**Figure 3.**
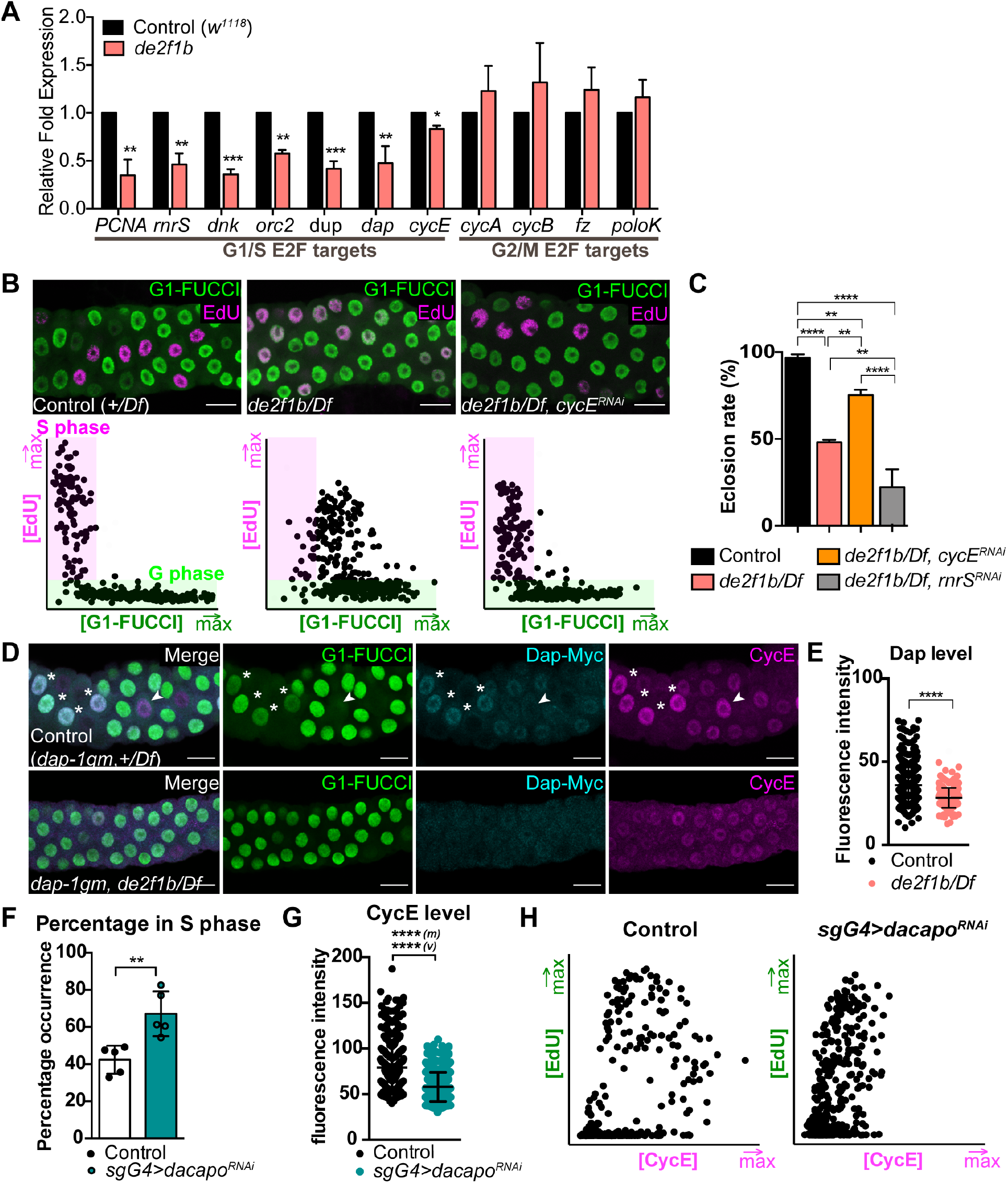
E2F1b regulates a specific subset of E2F targets that results in CycE hyperactivation. (A) RT-qPCR of E2F targets in control and *de2f1b*. Error bars indicate s.d. from three independent experiments. * = p ≤ 0.05; ** = p ≤ 0.01; *** =p ≤ 0.001. (B) Confocal micrographs of 80-85 hr AEL salivary glands labelled with G1-FUCCI and EdU. Salivary glands of control, *de2f1b,* and *de2f1b* with *cycE* depletion are presented. 2D scatter plots (lower panels) show their relative expression patterns. (C) Eclosion rates in control, *de2f1b,* and *de2f1b* expressing RNAi constructs against *cycE or rnrS* are compared. Error bars indicate s.d. from three independent experiments. ** = p ≤ 0.01; **** = p ≤ 0.0001. (D) Confocal micrographs of 80-85 hr AEL salivary glands expressing Myc-tagged Dap genomic construct (*dap-1gm*). Dap-Myc expression in control and *de2f1b* salivary glands is compared to G1-FUCCI and CycE expression. Asterisks indicate cells high in Dap-Myc, G1-FUCCI, and CycE. Arrowheads indicate cells absent of Dap-Myc and G1-FUCCI while CycE is still high. (E) A scatter plot showing differential Dap-Myc protein levels from control and *de2f1b* salivary glands. **** = p ≤ 0.0001. (F − H) The effect of depleting *dap* (*sgG4>dacapo^RNAi^*) is determined. (F) The percentage of EdU positive cells per salivary gland. Values represent the mean of five independent salivary glands and error bars indicate s.d. ** = p ≤ 0.01. (G) A scatter plot depicting CycE protein intensity from individual nuclei showing significant difference in (m) and (v). **** = p ≤ 0.0001. (H) 2D scatter plots showing the shift in the distribution of the relative expression pattern of EdU (green) and CycE (magenta) in upon *dap* depletion. All scale bars = 25 μm.

Importantly, there were notable discrepancies within the G1/S targets. While the expression of many G1/S targets, such as *rnrS* and *PCNA,* had an approximately 2-fold decrease, *cycE* levels are relatively unaffected in *de2f1b* salivary glands, resulting in less than 20% reduction. Interestingly, *dap*, a p27^CIP/KIP^-like Cdk inhibitor of CycE/Cdk2 activity [23], also showed a 2-fold decrease in *de2f1b* salivary glands. This is likely a tissue-specific effect since in *de2f1* mutant embryos, *dap* expression was shown to be independent of E2F1 [24]. These quantitative results were further validated through *in situ* hybridization and analysis of reporter constructs (Figure S1D). Overall, our data suggest that *cycE* expression can be compensated by the presence of E2F1a while expression of other E2F1 targets, such as *PCNA* and *rnrS,* cannot be compensated.

We next asked whether the disproportional change between *cycE* expression and other E2F1 target genes have functional consequences. We were particularly interested by the fact that *dap* expression is more reduced in *de2f1b* salivary glands than *cycE.* This raises the possibility that despite CycE not reaching high levels (Figure 1E), CycE/Cdk2 activity may be elevated due to a dampened expression of its inhibitor. Therefore, we examined the effect of reducing *cycE* levels in *de2f1b* salivary gland cells. Upon depletion of *cycE* using salivary gland specific driver *sgG4*, we observed a strong suppression of the cell cycle defect described in Figure 2D. *cycE* depletion in *de2f1b* salivary glands eliminates the population of cell where G1-FUCCI expression and EdU incorporation overlap (Figure 3B). This result suggests that this overlap is likely due to hyperactivation of CycE that pushes cells to prematurely enter S phase in *de2f1b* salivary glands. This is also supported by partial restoration of nuclear accumulation of DUP upon *cycE* depletion (Figure S1E, S1F). Furthermore, to determine if increased *cycE* activity contributes to the previously reported semi-lethality of *de2f1b* flies [9], *cycE* was weakly depleted in most fly tissues using *heatshock*-GAL4 (*hsG4*) without heat shock. Importantly, depleting *cycE* significantly rescued the pupal lethality to 75% eclosion rate from 50% while depleting *rnrS*, another well-known E2F1 target, increased the pupal lethality to 22% (Figure 3C). Taken together, our results indicate that hyperactive CycE is a major contributor of the defects observed in *de2f1b* flies.

### Dap sets the CycE/Cdk2 activity threshold for timely S phase entry

To determine the contribution of Dap to CycE hyperactivation, we first determined the if nuclear level of Dap is affected in *de2f1b* salivary glands. Because our commercially purchased Dap antibody did not result in direct detection of the endogenous protein levels, we used a Dap protein reporter (*dap-1gm*). *dap-1gm* is a transgene containing a 6-kb 5’ regulatory region of Dap with the coding sequence tagged with 6XMyc epitopes on the C-terminus [25]. *dap-1gm* faithfully recapitulates the endogenous Dap expression in most tissues with the exception of the extraembryonic amnioserosa and importantly rescues the semi-lethality of *dap* homozygous mutants [25]. Upon analyzing the expression of Dap (Dap-Myc) relative to G1-FUCCI and CycE (Figure 3D), we observed that Dap expression peaks in CycE expressing cells of control salivary glands only when they are in G phase (Figure 3D asterisks). In CycE high and G1-FUCCI negative S phase cells, Dap expression is almost undetectable and therefore no longer coupled to CycE (Figure 3D arrowhead). This is likely due to the PIP box-dependent degradation of Dap, by CRL4^Cdt2^ in S phase [26]. This result suggests that Dap is expressed in CycE high G phase cells to limit its activity prior to entering S phase. In *de2f1b* salivary glands, however, Dap expression is significantly reduced, reflecting the decrease in the transcription level (Figure 3D bottom panel and Figure 3E). This indicates that the failure to express Dap likely promotes hyperactivation of CycE in G phase and premature entry into S phase.

Importantly, it has been suggested that Dap does not contribute to the periodic silencing of CycE/Cdk2 during endocycles [13]. However, this was largely based on the lack of ploidy and size defects in *dap* mutant nuclei. Therefore, we knocked down *dap* and determined its effect in the cell cycle and CycE expression. Consistent with previous findings, depletion of *dap* does not affect the overall ploidy during endocycles ([13], Figure S2B). However, we observed a significant increase of S phase population, similar to what was observed in *de2f1b* salivary glands (Figure 3F). Moreover, CycE levels were significantly decreased in *dap* knockdown salivary glands (Figure S2A and 3G,H). This is likely due to a decrease in the threshold of CycE levels to reach the CycE/CdK2 activity that is required for S phase entry in the absence of its negative regulator Dap. This also provides an explanation in which *de2f1b* cells prematurely enter S phase without reaching high levels of CycE. Taken together, these results indicate that Dap is required during the endocycle to control CycE/Cdk2 activity prior to entering S phase, and that decrease level of Dap contributes to many aspects of the endocycle defects observed in *de2f1b* salivary glands.

### E2F1b-dependent transcription of PCNA controls PIP box-mediated degradation and establishes a negative feedback loop

Our findings indicate that PCNA is one of the E2F1b-specific targets (Figure 3A). As mentioned above, the degradation mechanism of E2F1 requires S phase-coupled chromatin-bound PCNA-dependent action of CRL4^Cdt2^. We hypothesized that PCNA contributes to the oscillation defect shown in Figure 1A, raising a possibility that E2F1b activity is directly linked to its own degradation, which is critical for the oscillatory nature of E2F1 expression during endocycles. Correspondingly, nuclear PCNA protein expression oscillates in control salivary glands and is significantly decreased in *de2f1b* salivary glands (Figure 4A,B). In control salivary glands, cells with high PCNA levels have low E2F1 expression (Figure 4A asterisks) and low PCNA level have high E2F1 expression (Figure 4A arrowhead). In *de2f1b* salivary glands, cells with high PCNA levels are absent, indicating that E2F1b is required for proper PCNA expression during S phase. To test the role of PCNA in establishing the biphasic oscillation of E2F1 and CycE, we depleted PCNA in the salivary gland and generated a plot profile of E2F1 and CycE expression (Figure 4C). Strikingly, PCNA depletion led to a complete loss of oscillations between E2F1 and CycE similar what was observed in *de2f1b* salivary glands and stabilized the expression of both proteins in virtually every cell. Supporting the notion that this effect is mainly through deregulating the PIP-dependent degradation program, PCNA depletion completely disturbed the G1-FUCCI oscillation as well (Figure S2C).

**Figure 4.**
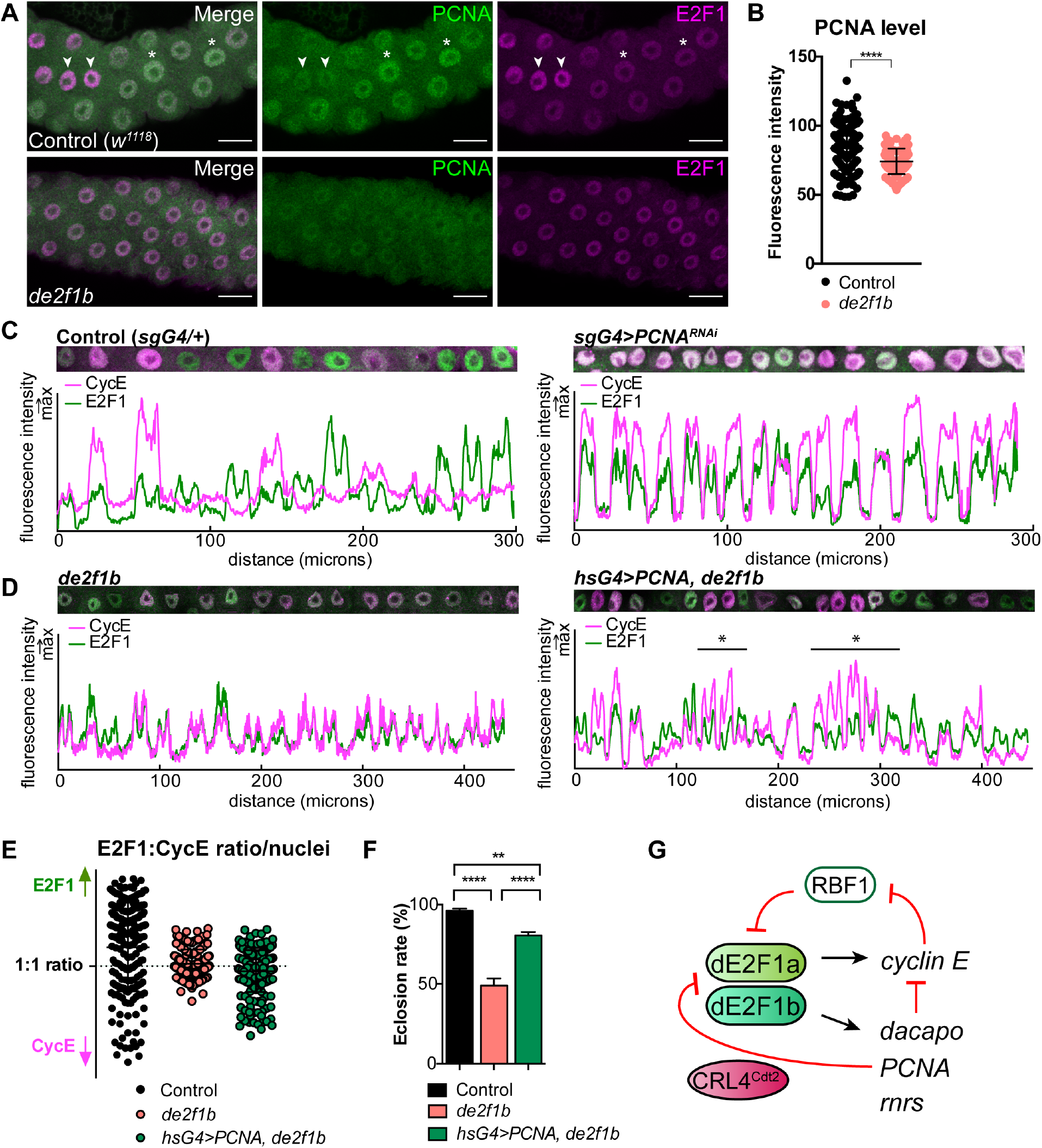
E2F1b-dependent transcription of PCNA controls PIP box-mediated degradation and establishes a negative feedback loop. (A) Confocal micrographs of 80-85 hr AEL Control and *de2f1b* salivary glands marking PCNA (green) and E2F1 (magenta). Arrowhead indicates high E2F1 and low PCNA cells. Asterisks indicate low E2F1 and high PCNA cells. Scale bars = 25 μm. (B) A scatter plot comparing the differential nuclear levels of PCNA in control and *de2f1b* salivary glands. **** = p ≤ 0.0001. (C) Plot profiles depicting the disruption of oscillation of E2F1 (green) and CycE (magenta) upon PCNA depletion by *sgG4>PCNA^RNAi^* from 80-85 hr AEL salivary glands. Plot profiles showing partial restoration of disrupted oscillation in *de2f1b* salivary glands of E2F1 (green) and CycE (magenta) through expression of PCNA (*hsG4>PCNA, de2f1b*). (E) E2F1:CycE Ratio plot is shown for the indicated genotypes showing partial restoration of high CycE cells upon PCNA expression in *de2f1b* salivary glands. (F) Eclosion rate in the described genotypes where PCNA expression significantly rescues the eclosion defect of *de2f1b* flies. ** = p ≤ 0.01; **** = p ≤ 0.0001. (G) A model illustrating the genetic circuitry controlled by E2F1a and E2F1b that is essential in endocycling cells.

Next, we determined the effect of partially restored PCNA expression in *de2f1b* salivary glands by weakly expressing PCNA under *hsG4* without heat shock. Strikingly, this resulted in cells where CycE and E2F1 is no longer coordinated (Figure 4D asterisk region). The restoration of CycE oscillation is best illustrated upon analysis of E2F1 and CycE intensity ratio in individual nuclei of the salivary gland (Figure 4E). In control cells, a wide range of [high E2F1]:[low CycE] to [low E2F1]:[high CycE] nuclei are observed. In *de2f1b* cells, most nuclei are close to the 1:1 ratio line, indicating that many nuclei have proportional levels of E2F1 and CycE. Partial restoration of PCNA expression in *de2f1b* cells notably changes the ratio profile whereby more [low E2F1]:[high CycE] cells are observed. Analysis of later staged salivary glands (105-110 hr AEL) revealed that that PCNA expression can improve the final ploidy of *de2f1b* salivary glands, making it comparable to the control (Figure S2D). However, we still observed the variable nuclear size defect seen in *de2f1b* salivary glands, suggesting that the nuclear size is independently regulated from ploidy control (Figure S2E, [9]). Importantly, weakly expressing PCNA in *de2f1b* flies significantly restored the eclosion rate (Figure 4F), similar to what is observed with depletion of *cycE* (Figure 3C). Taken together, our data suggest that the E2F1b-PCNA regulatory axis is crucial for proper degradation of E2F1 in S phase, which is required for downregulating *cycE* expression at the end of S phase.

### Coordinating the endocycle requires E2F1b-dependent genetic circuitry

Previous genetic studies have established E2F as an important regulator of the endocycle. The final ploidy of endocycling tissues is determined by silencing the E2F-dependent mitotic transcriptional program in mice and the regulation of the biphasic E2F1-CycE oscillators in flies [11]. In this report, we further refined the role of E2F during endocycle progression in Drosophila by identifying feedback mechanisms that govern the entry to and exit from S phase (Figure 4G). Endocycles consist of G and S phases, doubling DNA content without cell division. As a consequence, key cell cycle-dependent molecular events depend on controlling the timing of DNA synthesis. This includes proper licensing of origin of replication during G phase, firing of the origin upon entry to S phase, and preventing relicensing origin during S phase [27]. CycE/Cdk2 is at the center of these events. In G phase, activity of CycE/Cdk2 is held in check to ensure loading of licensing factors such as DUP/Cdt1. In contrast, CycE/Cdk2 becomes fully active during S phase to fire origins of replication and to prevent relicensing of origins. It has been previously thought that the nature of CycE/Cdk2 activity is directly contributed by the oscillating E2F1 expression and mainly regulated at the level of transcription in a cell cycle-dependent manner [13]. However, our data demonstrates that Dap is an integral part of the endocycle in salivary glands, fine tuning the timing of S phase entry. Curiously, Dap depletion does not result in ploidy defects in the salivary gland (Figure S2B and [13]). This is perhaps due to compensation by E2F1. Increased and/or premature CycE/Cdk2 activity likely inactivates RBF1 more efficiently, which in turn enables rapid accumulation of E2F1 targets such as ORCs, MCMs and DNA polymerases (Figure 4G). This would ensure expression of sufficient factors for DNA synthesis even if cells prematurely enter S phase. Nevertheless, an important implication of our finding is that the threshold of CycE/Cdk2 activity that is needed for entry to S phase may be regulated at two different levels: E2F-dependent transcriptional activation of *cycE* and Dap-mediated inhibition of CycE/Cdk2 activity.

By regulating the S phase-dependent expression of PCNA, E2F1b establishes a negative feedback loop to control its own degradation. In *de2f1b* salivary glands, cells have decreased PCNA expression, however, cells readily enter S phase due to hyperactive CycE/Cdk2. As a consequence, E2F1 destruction is no longer coupled to DNA synthesis and CycE is poorly downregulated at the end of S phase. This results in a deregulated endocycle in which E2F1 and CycE oscillation is lost (Figure 1A). Despite the loss of E2F1 and CycE oscillation, about 50% of *de2f1b* mutant complete morphogenesis and become adult flies (Figure 2C, 4F). This suggests that while E2F1 and CycE oscillation is an important regulatory feature, it is not absolutely essential for the endocycle as cells are able to become polyploid. To understand the mechanistic role of E2F-dependent target gene oscillations during the endocycle, it will be crucial to determine the consequences of *e2f* mutations on specific features of the endocycle, such as differential DNA replication, cell size and ploidy.

## Materials and Methods

### Fly strains

All fly strains and crosses were maintained at 25°C on standard cornmeal medium in a 12-hour light and dark cycle. Full list of stocks and genotypes for each experiment are described in Appendix Table S1 and S2.

### Eclosion Rate Quantification

To quantify the eclosion rate, eggs were collected from each genotype for 1 day then total pupae that eclosed after 14 days AEL were counted and their eclosion rate was calculated accordingly in comparison to total pupae present. Values represent an average of experimental triplicates. Error bars indicate standard deviation (s.d.).

### Immunostaining and EdU labelling

For immunostaining, 80-85 hr AEL salivary glands were fixed in 4% formaldehyde in 1XPBS for 20 minutes at room temperature (with the exception of dE2F1 immunostaining, where tissues were fixed at 4°C for 30 minutes). Tissues were then washed three times with 0.3% PBSTriton-X then three times with 0.1%PBSTriton-X. Tissues were incubated in 1% BSA blocking solution with appropriate primary antibody overnight at 4°C then washed three times in 0.1%PBSTriton-X. The secondary antibody incuation was carried out for 3 hours at room temperature, then tissues were washed four times in 0.1%PBSTriton-X and mounted for imaging. The following primary antibodies were used in this study: rabbit anti-dE2F1 (1:100, gift from N. Dyson, Massachusetts General Hospital), mouse anti-Myc (1:200, Developmental Studies Hybridoma Bank (DSHB) 9E10), guinea pig anti-DUP (1:1000, gift from T. Orr-Weaver and J. Nordman), goat anti-CycE (1:200, Santa Cruz sc15903), rabbit anti-CycE (1:100, Santa Cruz sc33748), mouse anti-PCNA (1:200, Cell Signaling PC10 #2586), and goat anti-GFP conjugated to FITC (1:100, Abcam ab6662). The following secondary antibodies coupled to fluorescent dyes from Jackson ImmunoResearch were used in 1:200 dilutions: donkey anti-rabbit cy2, donkey anti-goat cy3, donkey anti-goat cy5, goat anti-mouse cy3, donkey anti-mouse cy5, goat anti-guinea pig cy3. Ethynyl-2’-Deoxyuridine (EdU) cell proliferation assay (Invitrogen C10339) was used according to the manufacturer’s specifications. DNA was visualized with 0.1 μg/mL DAPI. Representative images were selected from a minimum of 10 independent tissues that were appropriately labeled.

### Microscopy

All fluorescently labelled tissues were mounted using a glycerol-based anti-fade mounting medium containing 5% N-propyl gallate and 90% glycerol in 1XPBS. All confocal micrographs were acquired using a laser-scanning Leica SP8 confocal microscope with a 20x/0.7 dry objective as a Z-stack with 1.5 μm set as the Z distance at the Cell Imaging and Analysis Network, McGill University. Representative images are individual slices from Z-stacks and a minimum of 10 independent confocal micrographs were taken. Images from *in situ* hybridization experiments were taken using the Canon Powershot G10 and Zeiss SteREO Discovery V8 modular stereo microscope with a conversion lens adaptor. All images were processed using Fiji (http://fiji.sc/Fiji).

### Generation of Plot Profile plots

Per genotype, a minimum of 250 microns was spanned across a section of the salivary gland through the middle of each nuclei using the freehand line tool in Fiji. The Plot Profile analysis tool was used to generate plots per individual channel. Generated plot profiles for two channels were merged on GraphPad Prism. The linearized image was created using the Straighten edit tool from Fiji.

### Quantification of nuclear protein and DAPI intensities

Nuclear protein intensities for dE2F1, CycE, G1-FUCCI, EdU, DUP, Dap, PCNA, Dap-Myc, and DAPI were obtained using the particle analysis tool in Fiji. For each salivary gland, a Z-projection of 2 stacks from the most apical and most basal sections were produced then subjected to analysis to accurately quantify the overall distribution of nuclear concentrations in a given salivary gland. Five independent salivary glands from the same experiment were analyzed to control for differential protein intensities. A minimum of 300 nuclei are shown for all analyses. Nuclear protein intensities were then plotted as a X, Y point plot or a scatter plot to show protein levels. Ploidy was plotted in a box and whiskers plot.

### Quantification of S phase population

EdU incorporation positive nuclei were identified by the particle analysis tool in Fiji. For each salivary gland, the percentage of EdU incorporation positive nuclei per salivary gland was counted for five independent salivary glands from Z-projection of 2 stacks from the most apical and most basal sections. The average of the S phase populations taken from five salivary glands is shown. Error bars indicate s.d.

### Quantification of E2F1:CycE ratio

Fluorescence intensities of E2F1 and CycE were determined using the particle analysis tool in Fiji. For each nucleus, E2F1 intensity was divided by CycE intensity then log transformed (log base 10) to get values ranging from 1 to −1, where 0 represents the 1:1 ratio (labelled as dotted line), values towards 1 represent cells with higher E2F1, and values towards −1 represent cells with higher CycE.

### Quantification of relative standard deviation of nuclear area

Nuclear area was determined by using the particle analysis tool in Fiji. The relative standard deviation was calculated by dividing the standard deviation of nuclear size by the mean of the nuclear size per salivary gland. Three independent salivary glands were analyzed for each genotype. Error bars indicate s.d.

### RNA extraction and cDNA synthesis

RNA was extracted from 40 *w*^*1118*^ or *de2f1b* 80-85 hr AEL salivary glands using the Aurum Total RNA Mini Kit (Bio-Rad) with treatment of DNase I to remove genomic DNA contamination. 200 ng of RNA was used to synthesize cDNA using the iScript cDNA Synthesis Kit (Bio-Rad).

### Reverse Transcriptase quantitative PCR (RT-qPCR)

Gene expression was quantified using the DyNAmo Flash SYBR Green qPCR Kit (Thermo-Scientific) with the comparative cycle method using the Bio-Rad CFX 96 Real-Time System and C1000 Thermal Cycler. Two housekeeping genes, *rp49* and β*-tubulin*, were used to normalize the data obtained. All values represent averages of experimental biological triplicates and error bars represent s.d. All primers were designed using Primer3 (Whitehead Institute for Biomedical Research, http://Frodo.wi.mit.edu/primer3/ or taken from the DRSC/TRiP Functional Genomics Resources FlyPrimerBank [28]. All primers were tested and run on an 8% acrylamide gel to ensure a single PCR product. Primers used in this paper are listed on Appendix Table S3.

### Statistical analysis

Statistical analyses were performed using the GraphPad Prism software. Two-tailed unpaired t-tests were performed for Figures 1D, 1E, 3A, 3E, 3F, 3G, 4B, and S1A. F-test was performed to calculate the variance for Figures 1G and 3G. One-way ANOVA was performed on Figures 3C, 4F, S1F, S2B, S2E, and S2F. P-values represent ns = p > 0.05; * = p ≤ 0.05; ** = p ≤ 0.01; *** =p ≤ 0.001; **** = p ≤ 0.0001.

## Acknowledgements

We are grateful to T. Orr-Weaver (Whitehead Institute, MIT), J. Nordman (Vanderbilt University), and C. Lehner (University of Zurich) for sharing antibodies and fly stocks. We thank the Bloomington Stock Center, Drosophila Genetic Resource Center, and Vienna Drosophila Research Center for providing fly stocks. We also thank CIAN/ABIF for their assistance in confocal image acquisition.

## Supplemental Materials

**Supplemental Figure 1.**
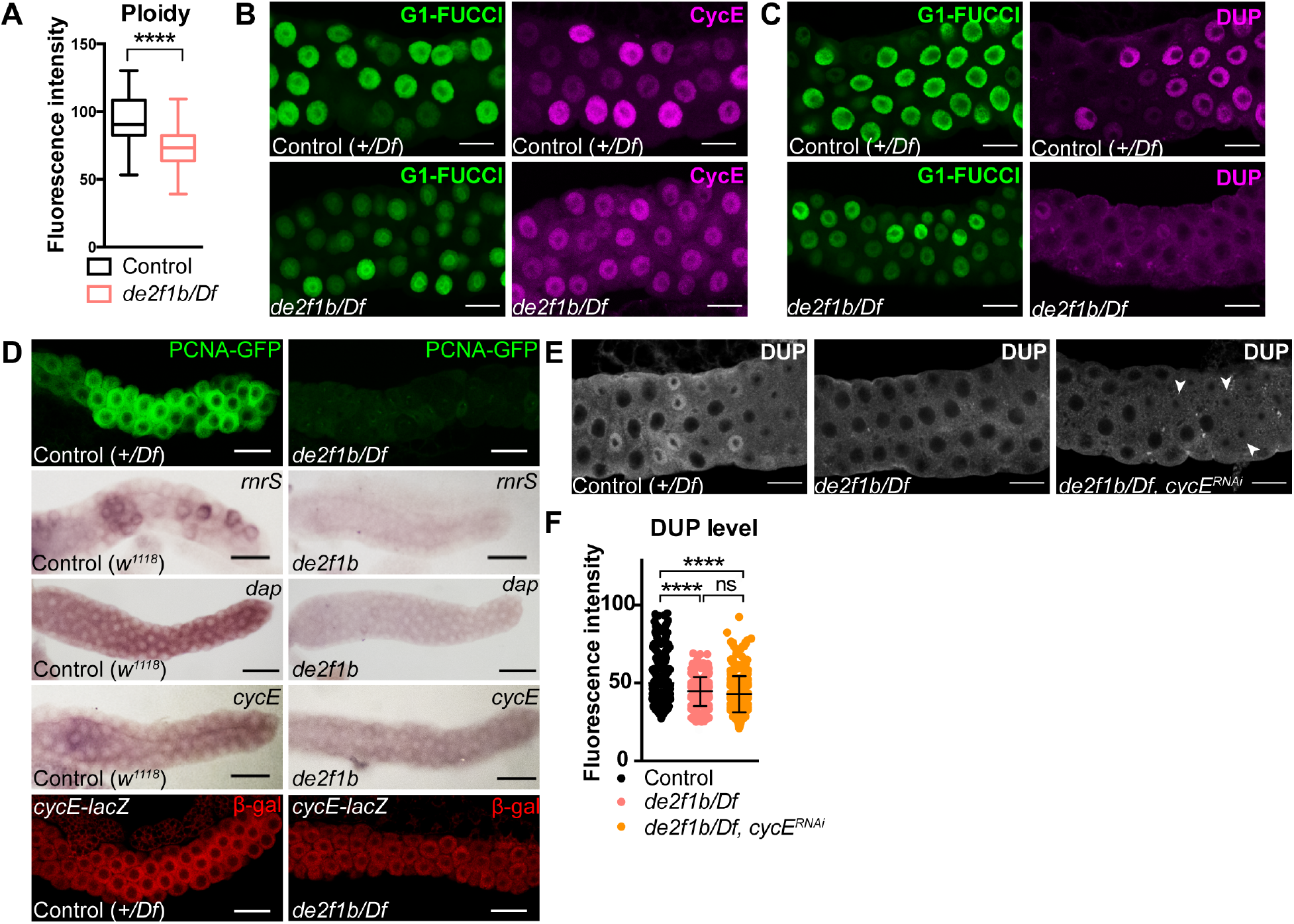
Analysis of *de2f1b* salivary glands. (A) A box and whiskers plot showing differences in ploidy in control and *de2f1b/Df* salivary glands. **** = p ≤ 0.0001. (B, C) Confocal micrograph of 80-85 hr AEL salivary glands of control and *de2f1b/Df* showing individual channel images of G1-FUCCI (green) and CycE (magenta) from Figure 2E and G1-FUCCI (green) and DUP (magenta) from Figure 2F, respectively. (D) Micrographs of 80-85 hours AEL control and *de2f1b* or *de2f1b/Df* salivary glands. The expression patterns of PCNA-GFP reporter, which is derived from the PCNA promoter and *rnrS*, *dap*, *cycE* transcripts and *cycE-lacZ* reporter are shown. (E) Confocal micrographs of 80-85 hr AEL salivary glands from indicated genotypes showing partial restoration of nuclear DAP upon *cycE* depletion in *de2f1b* salivary glands (arrowheads). (F) Quantification of DUP levels from salivary glands shown in (E). ns = p > 0.05; **** = p ≤ 0.0001. All scale bars = 25 μm.

**Supplemental Figure 2.**
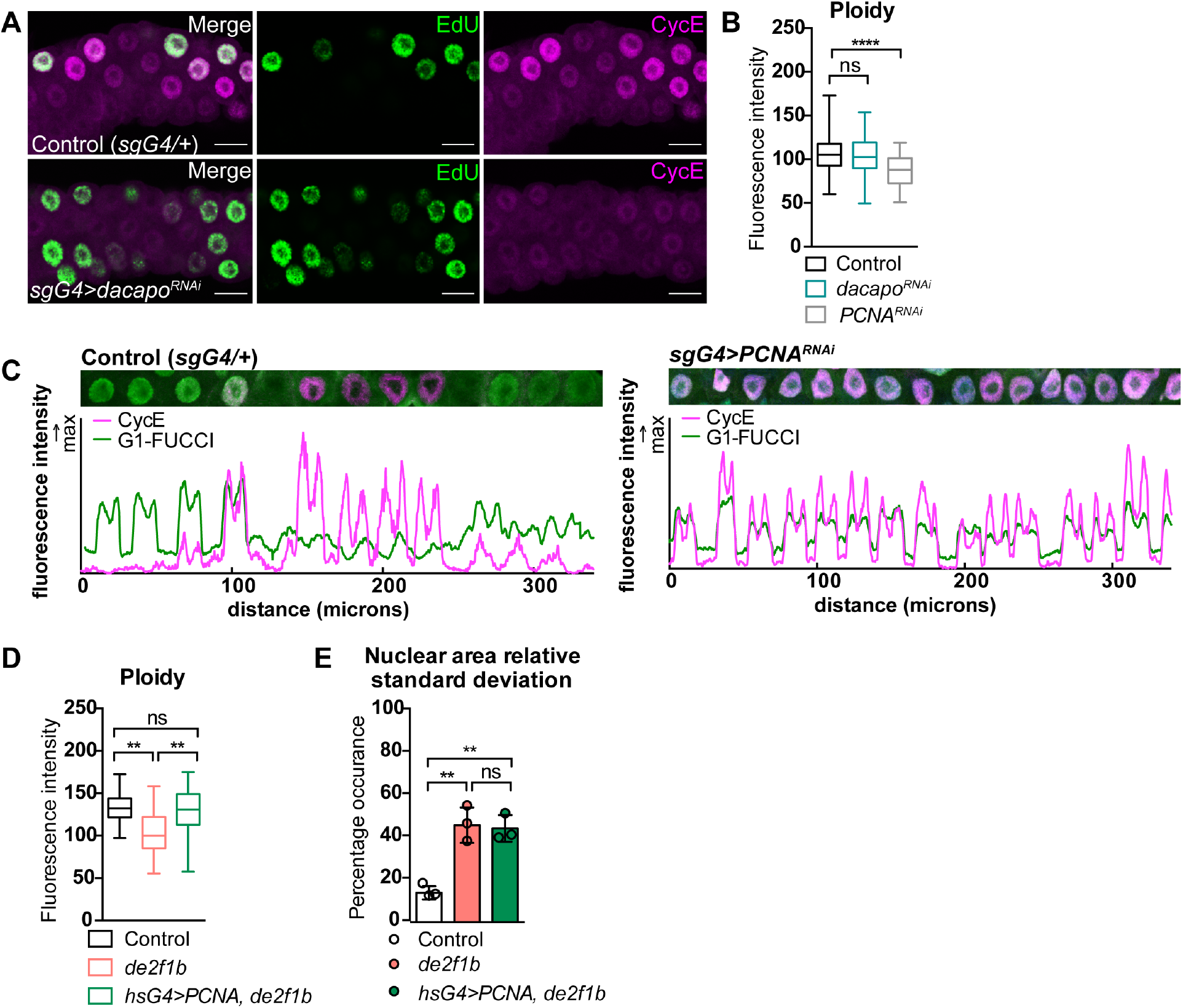
Dap and PCNA are crucial E2F1b targets during the endocycle. (A) Confocal micrographs of 80-85 hr AEL salivary glands labelled with EdU and CycE in control versus *dap* depletion (*sgG4>dap^RNAi^*). Scale bars = 25 μm. (B) Ploidy quantification between control, *sgG4>dacapo^RNAi^*, and *sgG4>PCNA^RNAi^* in 80-85 hr AEL salivary glands. ns = p > 0.05; **** = p ≤ 0.0001. (C) Plot profiles depicting nuclear intensities of G1-FUCCI (green) and CycE (magenta) from 80-85 hr AEL salivary gland. The effect of depleting PCNA *sgG4>PCNA^RNAi^* is shown. (D) Ploidy quantification in 105-110 hr AEL salivary glands of the indicated genotypes. Expressing PCNA in *de2f1b* salivary glands rescue the ploidy defect. ns = p > 0.05; ** = p ≤ 0.01. (E) Average of the relative standard deviation of nuclear area of indicated genotypes showing expressing PCNA in *de2f1b* salivary glands do not rescue the highly variable nuclear size. Error bars indicate s.d. ns = p > 0.05; ** = p ≤ 0.01.

**Appendix Table S1.** Drosophila stocks used in this study

| Stock | Source |
| --- | --- |
| *w<sup>1118</sup>* | Bloomington Drosophila Stock Center (BDSC) 5905 |
| *sgs3-GAL4* | BDSC 6870 |
| *heatshock-GAL4* | BDSC 2077 |
| *UAS-cycE<sup>RNAi</sup>* | BDSC 36092 |
| *UAS-rnrS<sup>RNAi</sup>* | BDSC 32864 |
| *UAS-dacapo<sup>RNAi</sup>* | BDSC 64026 |
| *UAS-PCNA<sup>RNAi</sup>* | Vienna Drosophila Resource Center (VDRC) GD51253 |
| *UAS-GFP-E2F1<sup>1-230</sup>, UAS-mRFP<sup>nls</sup>-CycB<sup>1-266</sup>* | BDSC 55121 |
| *cycE-lacZ<sup>16.4kb</sup>* | BDSC 30722 |
| *Df(3R)Exel6186* | BDSC 7665 |
| *dap-1gm* | Provided by C. Lehner [25] |
| *3XPCNA-GFP* | Provided by R. Duronio [29] |
| *UAS-PCNA* | This study, generated using the Drosophila Gateway collection (Drosophila Genomic Resource Center) |
| *de2f1b* | Lab collection [9] |
| *Df(3R)Exel6186, UAS-rnrS<sup>RNAi</sup>* | Lab collection<br>Recombined lines |
| *Df(3R)Exel6186, 3XPCNAGFP* | Lab collection<br>Recombined lines |
| *Df(3R)Exel6186, sgG4* | Lab collection<br>Recombined lines |
| *Df(3R)Exel6186, cycE-lacZ<sup>16.4kb</sup>* | Lab collection<br>Recombined lines |

**Appendix Table S2.** Description of full genotypes for each experiment performed in this study.

| Figure | Genotype |
| --- | --- |
| 1A-F,<br>S1A | **Control:**<br>*w;; +/Df(3R)Exel6186*<br>**de2f1b/Df:**<br>*w;; de2f1b/Df(3R)Exel6186* |
| 2C-F,<br>S1B,C | **Control:**<br>*w; +/UAS-GFP-E2F1<sub>1-230</sub>, UAS-mRFP<sup>nls</sup>-CycB<sub>1-266</sub>; +/Df(3R)Exel6186, sgG4*<br>**de2f1b/Df:**<br>*w; +/UAS-GFP-E2F1<sub>1-230</sub>, UAS-mRFP<sup>nls</sup>-CycB<sub>1-266</sub>; de2f1b/Df(3R)Exel6186, sgG4* |
| 3B<br>S1E,F | **Control:**<br>*w; +/UAS-GFP-E2F1<sub>1-230</sub>, UAS-mRFP<sup>nls</sup>-CycB<sub>1-266</sub>; +/Df(3R)Exel6186, sgG4*<br>**de2f1b/Df:**<br>*w; +/UAS-GFP-E2F1<sub>1-230</sub>, UAS-mRFP<sup>nls</sup>-CycB<sub>1-266</sub>; de2f1b/Df(3R)Exel6186, sgG4*<br>**de2f1b/Df, cycE<sup>RNAi</sup>:**<br>*w; UAS-cycE<sup>RNAi</sup>/UAS-GFP-E2F1<sub>1-230</sub>, UAS-mRFP<sup>nls</sup>-CycB<sub>1-266</sub>; de2f1b/Df(3R)Exel6186, sgG4* |
| 3C | **Control:**<br>*w;; +/Df(3R)Exel6186)*<br>**de2f1b/Df:**<br>*w;; de2f1b/Df (3R)Exel6186*<br>**de2f1b/Df, cycE<sup>RNAi</sup>:**<br>*w; hsG4/UAS-cycE<sup>RNAi</sup>; de2f1b/Df(3R)Exel6186*<br>**de2f1b/Df, rnrS<sup>RNAi</sup>:**<br>*w; hsG4/+; de2f1b/Df(3R)Exel6186, UAS-rnrS<sup>RNAi</sup>.* |
| 3D,E | **Control (dap-1gm, +/Df):**<br>*w; dap-1gm/UAS-GFP-E2F1<sub>1-230</sub>, UAS-mRFP<sup>nls</sup>-CycB<sub>1-266</sub>; +/Df(3R)Exel6186, sgG4*<br>**dap-1gm, de2f1b/Df:**<br>*w; dap-1gm/UAS-GFP-E2F1<sub>1-230</sub>, UAS-mRFP<sup>nls</sup>-CycB<sub>1-266</sub>; de2f1b/Df(3R)Exel6186, sgG4* |
| 3F-H<br>S2A<br>S2B | **Control:**<br>*w;; +/sgG4*<br>**sgG4>dacapo<sup>RNAi</sup> / dacapo<sup>RNAi</sup>:**<br>*w; +/UAS-dacapo<sup>RNAi</sup>; +/sgG4* |
| 4C<br>S2B | **Control :**<br>*w;; +/sgG4*<br>**sgG4>PCNA<sup>RNAi</sup> / PCNA<sup>RNAi</sup>:**<br>*w; +/UAS-PCNA<sup>RNAi</sup>; +/sgG4* |
| 4D-F<br>S2D-F | **hsG4>PCNA; de2f1b:**<br>*w; hsG4/UAS-PCNA; de2f1b* |
| S1D | <u>First panel</u><br>**Control (+/Df):**<br>*w;; +/Df(3R)Exel6186, 3XPCNAGFP*<br>**de2f1b/Df:**<br>*w;; de2f1b/Df(3R)Exel6186, 3XPCNAGFP*<br><u>Fourth panel</u><br>**Control (+/Df):**<br>*w;; +/Df(3R)Exel6186, cycE-lacZ<sup>16.4kb</sup>*<br>**de2f1b/Df:**<br>*w;; de2f1b/Df(3R)Exel6186, cycE-lacZ<sup>16.4kb</sup>* |
| S2C | **Control :**<br>*w;+/UAS-GFP-E2F1<sub>1-230</sub>, UAS-mRFP<sup>nls</sup>-CycB<sub>1-266</sub> ; +/sgG4*<br>**sgG4>PCNA<sup>RNAi</sup> / PCNA<sup>RNAi</sup>:**<br>*w; UAS-GFP-E2F1<sub>1-230</sub>, UAS-mRFP<sup>nls</sup>-CycB<sub>1-266</sub>/UAS-PCNA<sup>RNAi</sup>;*<br>*+/sgG4* |

**Appendix Table S3.**
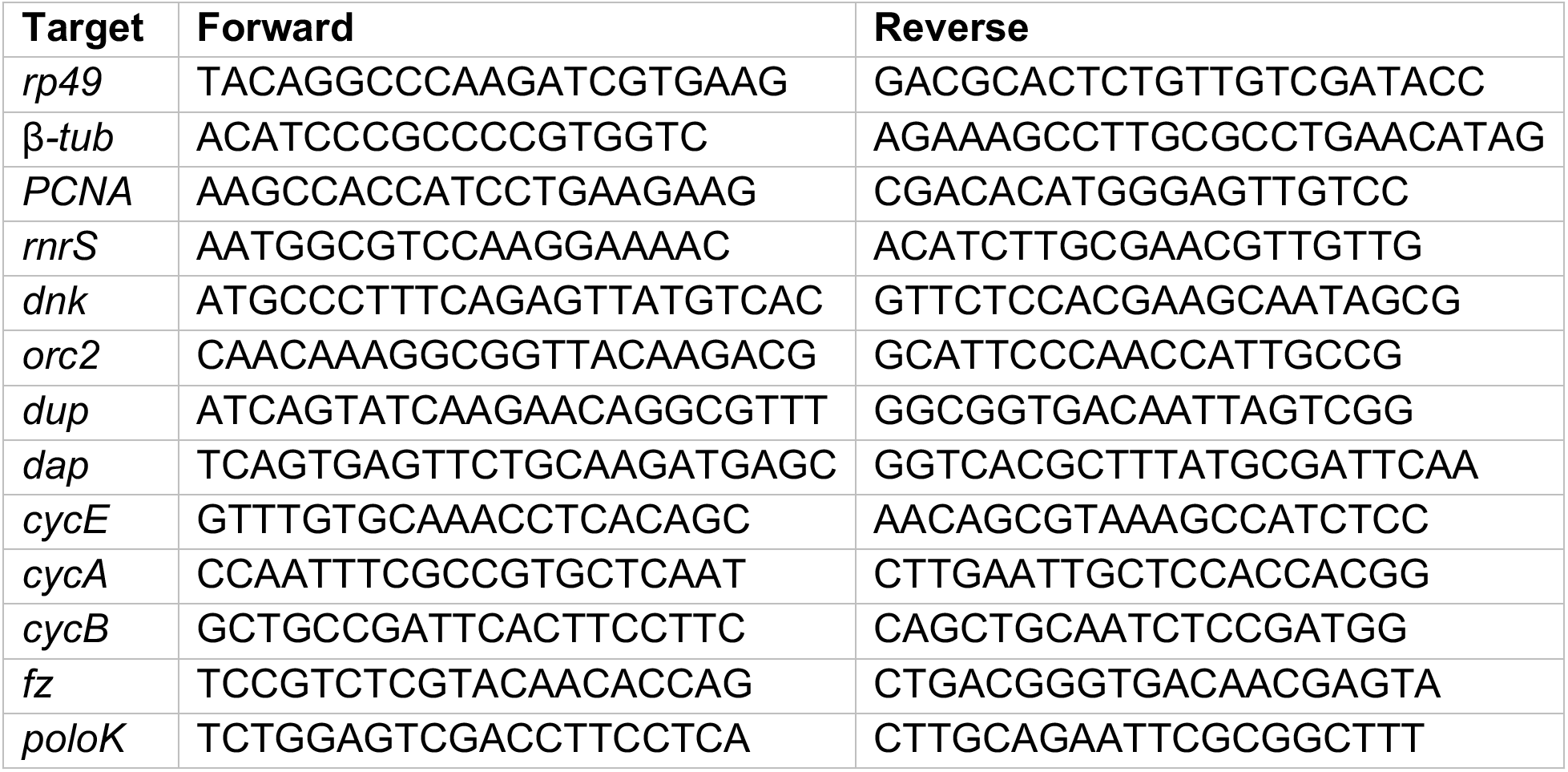
List of primers used in this study.

## Supplemental Materials and Methods

### *In situ* hybridization

80-85 hr AEL salivary glands were prepared for *in situ* hybridization as described previously [30]. Briefly, after hybridization of DIG-labeled probes, samples were incubated with Alkaline Phosphatase conjugated anti-DIG antibody (Roche) then subjected to detection using the NBT/BCIP stock solution (Roche). A minimum of 10 independent salivary glands were examined and representative images were selected. *rnrS*, *cycE* probes were generated as previously described [31] and the *dap* probe was prepared during this study (details below).

### *dacapo* RNA probe generation

*dap* probe was generated using standard PCR-based probe generation methods as described previously [32]. In brief, the *dap* coding sequence was flanked by T7 and T3 polymerase sequences in the forward and reverse primers, respectively, using the following primers:

Forward primer with T7:

5’-TAATACGACTCACTATAGGGAGAATGGTCAGTGCCCGAGTCCTGAATCC-3’

Reverse primer with T3:

5’-AATTAACCCTCACTAAAGGGAGATTAGTTGTGGCGCGGCCGCTTCAAC-3’ PCR was performed using standard procedures for the Phusion High Fidelity DNA polymerase (NEB) using the *dap* cDNA clone obtained from the Drosophila Genomics Resource Center (#RE12958). PCR product was purified then subjected to DIG-labeled transcription by T7 or T3 polymerase to generate the sense and anti-sense probes.

## References

1. Lilly MA, Duronio RJ (2005) New insights into cell cycle control from the Drosophila endocycle. Oncogene 24: 2765–2775.

2. Shu Z, Row S, Deng W-M (2018) Endoreplication: The Good, the Bad, and the Ugly. Trends Cell Biol.

3. Zielke N, Edgar BA, DePamphilis ML (2013) Endoreplication. Cold Spring Harb Perspect Biol 5: a012948.

4. Chen H-Z, Ouseph MM, Li J, Pécot T, Chokshi V, Kent L, Bae S, Byrne M, Duran C, Comstock G, et al. (2012) Canonical and atypical E2Fs regulate the mammalian endocycle. Nat Cell Biol 14: 1192–1202.

5. Pandit SK, Westendorp B, Nantasanti S, Van Liere E, Tooten PCJ, Cornelissen PWA, Toussaint MJM, Lamers WH, de Bruin A (2012) E2F8 is essential for polyploidization in mammalian cells. Nat Cell Biol 14: 1181–1191.

6. Weng L, Zhu C, Xu J, Du W (2003) Critical role of active repression by E2F and Rb proteins in endoreplication during Drosophila development. EMBO J 22: 3865–3875.

7. Royzman I, Austin RJ, Bosco G, Bell SP, Orr-Weaver TL (1999) ORC localization in Drosophila follicle cells and the effects of mutations in dE2F and dDP. Genes Dev 13: 827–840.

8. Royzman I, Hayashi-Hagihara A, Dej KJ, Bosco G, Lee JY, Orr-Weaver TL (2002) The E2F cell cycle regulator is required for Drosophila nurse cell DNA replication and apoptosis. Mech Dev 119: 225–237.

9. Kim M, Tang JP, Moon NS (2018) An alternatively spliced form affecting the Marked Box domain of Drosophila E2F1 is required for proper cell cycle regulation. PLoS Genet 14: e1007204.

10. Dimova DK, Dyson NJ (2005) The E2F transcriptional network: old acquaintances with new faces. Oncogene 24: 2810–2826.

11. Edgar BA, Zielke N, Gutierrez C (2014) Endocycles: a recurrent evolutionary innovation for post-mitotic cell growth. Nat Rev Mol Cell Biol 15: 197–210.

12. Shibutani ST, la Cruz de AFA, Tran V, Turbyfill WJ, Reis T, Edgar BA, Duronio RJ (2008) Intrinsic negative cell cycle regulation provided by PIP box- and Cul4Cdt2-mediated destruction of E2f1 during S phase. Developmental Cell 15: 890–900.

13. Zielke N, Kim KJ, Tran V, Shibutani ST, Bravo M-J, Nagarajan S, van Straaten M, Woods B, Dassow von G, Rottig C, et al. (2011) Control of Drosophila endocycles by E2F and CRL4(CDT2). Nature 480: 123–127.

14. Li J, Ran C, Li E, Gordon F, Comstock G, Siddiqui H, Cleghorn W,Chen H-Z, Kornacker K, Liu C-G, et al. (2008) Synergistic function of E2F7 and E2F8 is essential for cell survival and embryonic development. Developmental Cell 14: 62–75.

15. Moon NS, Dyson N (2008) E2F7 and E2F8 keep the E2F family in balance. Developmental Cell 14: 1–3.

16. Zielke N, Korzelius J, van Straaten M, Bender K, Schuhknecht GFP, Dutta D, Xiang J, Edgar BA (2014) Fly-FUCCI: A versatile tool for studying cell proliferation in complex tissues. Cell Rep 7: 588–598.

17. Shibutani ST, la Cruz de AFA, Tran V, Turbyfill WJ, Reis T, Edgar BA, Duronio RJ (2008) Intrinsic negative cell cycle regulation provided by PIP box- and Cul4Cdt2-mediated destruction of E2f1 during S phase. Developmental Cell 15: 890–900.

18. Whittaker AJ, Royzman I, Orr-Weaver TL (2000) Drosophila double parked: a conserved, essential replication protein that colocalizes with the origin recognition complex and links DNA replication with mitosis and the down-regulation of S phase transcripts. Genes Dev 14: 1765–1776.

19. Thomer M, May NR, Aggarwal BD, Kwok G, Calvi BR (2004) Drosophila double-parked is sufficient to induce re-replication during development and is regulated by cyclin E/CDK2. Development 131: 4807–4818.

20. Arias EE, Walter JC (2006) PCNA functions as a molecular platform to trigger Cdt1 destruction and prevent re-replication. Nat Cell Biol 8: 84–90.

21. Jin J, Arias EE, Chen J, Harper JW, Walter JC (2006) A Family of Diverse Cul4-Ddb1-Interacting Proteins Includes Cdt2, which Is Required for S Phase Destruction of the Replication Factor Cdt1. Mol Cell 23: 709–721.

22. Senga T, Sivaprasad U, Zhu W, Park JH, Arias EE, Walter JC, Dutta A (2006) PCNA is a cofactor for Cdt1 degradation by CUL4/DDB1-mediated N-terminal ubiquitination. J Biol Chem 281: 6246–6252.

23. de Nooij JC, Letendre MA, Hariharan IK (1996) A Cyclin-Dependent Kinase Inhibitor, Dacapo, Is Necessary for Timely Exit from the Cell Cycle during Drosophila Embryogenesis. Cell 87: 1237–1247.

24. de Nooij JC, Graber KH, Hariharan IK (2000) Expression of the cyclin-dependent kinase inhibitor Dacapo is regulated by Cyclin E. Mech Dev 97: 73–83.

25. Meyer CA, Kramer I, Dittrich R, Marzodko S, Emmerich J, Lehner CF (2002) Drosophila p27Dacapo expression during embryogenesis is controlled by a complex regulatory region independent of cell cycle progression. Development 129: 319–328.

26. Swanson CI, Meserve JH, McCarter PC, Thieme A, Mathew T, Elston TC, Duronio RJ (2015) Expression of an S phase-stabilized version of the CDK inhibitor Dacapo can alter endoreplication. Development 142: 4288–4298.

27. Zielke N, Edgar BA, DePamphilis ML (2013) Endoreplication. Cold Spring Harb Perspect Biol 5: a012948–a012948.

28. Hu Y, Sopko R, Foos M, Kelley C, Flockhart I, Ammeux N, Wang X, Perkins L, Perrimon N, Mohr SE (2013) FlyPrimerBank: an online database for Drosophila melanogaster gene expression analysis and knockdown evaluation of RNAi reagents. G3 (Bethesda) 3: 1607–1616.

29. Thacker SA, Bonnette PC, Duronio RJ (2003) The contribution of E2F-regulated transcription to Drosophila PCNA gene function. Curr Biol 13: 53–58.

30. Du W (2000) Suppression of the rbf null mutants by a de2f1 allele that lacks transactivation domain. Development 127: 367–379.

31. Hsieh T-C, Nicolay BN, Frolov MV, Moon NS (2010) Tuberous sclerosis complex 1 regulates dE2F1 expression during development and cooperates with RBF1 to control proliferation and survival. PLoS Genet 6: e1001071.

32. Legendre F, Cody N, Iampietro C, Bergalet J, Lefebvre FA, Moquin-Beaudry G, Zhang O, Wang X, Lécuyer E (2013) Whole mount RNA fluorescent in situ hybridization of Drosophila embryos. J Vis Exp e50057.

